# Group A streptococci induce high-affinity M protein-fibronectin interaction when specific human antibodies are bound

**DOI:** 10.1101/2022.05.04.490590

**Authors:** Sebastian Wrighton, Vibha Kumra Ahnlide, Oscar André, Wael Bahnan, Pontus Nordenfelt

## Abstract

Group A streptococcus (GAS) is a highly adapted, human-specific pathogen that is known to manipulate the immune system through various mechanisms. GAS’ M protein constitutes a primary target of the immune system due to its spatial configuration and dominance on the bacterial surface. Antibody responses targeting the M protein have been shown to favor the conserved C region. Such antibodies circumvent antigenic escape and efficiently bind to various M types. The ability of GAS to bind to fibronectin (Fn), a high molecular weight glycoprotein of the extracellular matrix, has long been known to be essential for the pathogen’s evolutionary success and fitness. However, some strains lack the ability to efficiently bind Fn. Instead, they have been found to inefficiently bind Fn via the M protein A-B domains. Here, we show that human antibodies can induce a high-affinity Fn-binding state in M proteins, likely by enhancing the weak A-B domain binding. The antibodies bind to a conserved region of M proteins, and the high-affinity binding only occurs on the individual M proteins with bound specific antibodies. By allowing the binding of antibodies to a certain region in M, and thereby enhancing Fn-binding, GAS exploits the human humoral immune response to efficiently bind Fn without needing to waste energy on the production of additional proteins – potentially giving such strains an evolutionary advantage.

## Introduction

*Streptococcus pyogenes*, also commonly referred to as group A streptococcus (GAS), is an extremely successful, ubiquitously found, human pathogen causing >600 million infections each year. While for the most part these infections result in mild disease development, more severe and invasive cases result in a mortality rate of up to 25% (1, 2). Fibronectin (Fn), a high-molecular-weight, glycoprotein of the extracellular matrix (ECM), is important due to its ability to serve as an adaptor protein of the ECM, allowing cells to interact with their environment. This is enabled by heterodimeric cell surface receptors known as integrins (3). Fn exists in an insoluble form whereby it is a constituent of many extracellular matrices and a soluble form that is found in various bodily fluids (3, 4). It is a protein dimer consisting of two almost identical monomers with multiple binding domains (5). The binding of Fn by GAS is a well-documented phenomenon most commonly implicated in enhancing adhesion to, and invasion of host cells (4, 6, 7). This binding is mediated by surface-expressed bacterial proteins which possess Fn-specific binding domains. Bound Fn can thus promote host tissue invasion by interacting with integrins expressed on epithelial cells (8–10). As many as 12 GAS surface proteins have been identified as directly or indirectly facilitating the binding of Fn (11). This fact alone hints at an evolutionarily driven process since bacteria would not waste energy on the production of superfluous proteins.

The GAS M protein exemplifies how the bacterium has evolved to manipulate the host immune system. Since the N-terminal domain is highly variable between various GAS serotypes it has become common practice to classify GAS depending on the specific M protein they express (12, 13). M protein dominates the surface of the bacterium and thus, is likely one of the main targets of the humoral immune response. Despite this, it has been shown that antibody responses to M only weakly target the more exposed variable regions and preferentially target the conserved C region (14, 15). Due to their conserved nature, these serve as cross-serotype antigens for the adaptive immune system (16). The successful development of a vaccine against GAS has eluded researchers for many years. This has mainly been due to the complex immune response to M protein. While it is the conserved regions that have been shown to elicit the more robust cross-species immune responses, it is these regions that have also been implicated in inducing various autoimmune sequelae (17, 18). Moreover, certain M types have been shown to possess the ability to bind Fn and thus increase adhesion or trigger internalization (19, 20). Specifically, the M1 protein has been studied in detail and while the exact binding mechanism remains unclear, it was possible to identify potential Fn-binding regions to the two apical domains (A and B) (21).

In recent years, advances in sequencing technology have uncovered a plethora of Fn-binding proteins being expressed by a wide range of bacterial species (22). It is generally accepted that bacteria primarily express proteins that improve or maintain their respective fitness. Since the cost of protein production is so immense, low-benefit proteins are not maintained and eventually, their encoding genes will be lost altogether (23, 24). This highlights the critical role that Fn-binding must play for GAS fitness and perhaps contributed to its success as a species. M1 GAS has for long periods been observed to be the most prevalent M strain – especially regarding strains implicated in invasive disease (12). Here, we show that certain M types, including M1, have evolved to exploit human antibodies targeting M. The antibodies target a conserved and immunodominant region in M protein, making them commonly available to the streptococci. The binding of these antibodies allows M protein to bind Fn with high affinity, making additional specialized proteins redundant. The M protein is well known to be versatile, and this adds a way in which group A streptococci can use the host’s immune response to its advantage and gain an evolutionary edge.

## Results

### Convalescent patient plasma enhances Fn-binding to GAS

Plasma contains large quantities of Fn (5) which is present in its various isoforms. It has been shown that the adaptive immune system will target surface-bound bacterial Fn-binding proteins with antibodies, leading to better vaccine provoked protection (25, 26). To assess whether GAS infections can lead to in vivo generation of antibodies that can neutralize Fn-binding proteins, we incubated the M1 serotype SF370, expressing GFP, in various dilutions of plasma-derived from convalescent donors who had recently recovered from a severe invasive GAS infection (Fig. 1a). As a control, we used plasma from healthy donors. After incubation, the bacteria were washed in PBS and the bound Fn was stained using an anti-human Fn primary antibody and a fluorescent anti-mouse secondary. Fibronectin bound to GAS was assessed by flow cytometry. Surprisingly, GAS incubated in the convalescent plasma consistently showed higher levels of bound Fn compared to the healthy controls. This was the case for all dilutions, whereby 50% plasma led to the most Fn being bound. This discrepancy was observed between all individuals except one healthy donor where the difference was not as pronounced. To determine whether the same would apply to plasma from a convalescent donor whose blood was used to derive anti-GAS monoclonals (27) we repeated the assay with plasma from this specific donor (Fig. 1b). For this experiment pooled plasma (from >20 healthy individuals) was used as a control and bacteria were left untreated in culture media to assess background autofluorescence. Fn-binding to GAS was measured by flow cytometry. As can be seen from the respective MFI values of the bound Fn, GAS incubated in dilutions of the antibody donor plasma consistently bound more Fn compared to those incubated in pooled plasma. For all dilutions, the untreated and pooled plasma showed the same binding level, whereas there was a dose-dependent increase when using donor plasma. In summary, incubating GAS in plasma derived from convalescent patients led to increased Fn-binding compared to when GAS was incubated in plasma from healthy donors. When repeating the assay using plasma from the original convalescent donor – from which our anti-GAS monoclonals are derived – we saw that plasma derived from the Ab donor was also able to promote increased Fn-binding compared to pooled plasma. **Certain monoclonal antibodies induce increased Fn-binding affinity to M1 GAS in an M protein-dependent manner**. Previous experiments (Fig 1a-b) showed that incubating GAS in plasma from GAS convalescent donors consistently results in increased Fn-binding. Next, we wanted to assess whether this was due to the antibodies contained within the respective plasma samples. We therefore treated SF370 GAS with fluorescently conjugated Fn isolated from plasma in conjunction with monoclonal antibodies (Figure 2a). The antibodies used as treatments were the monoclonals derived from the aforementioned convalescent donor (Ab25, Ab32, and Ab49). Furthermore, we used a non-binding control monoclonal (Xolair) and pooled intravenous antibodies (IVIG) as a negative and positive binding control respectively. GAS-bound fibronectin was assessed by flow cytometry. Treatment with the anti-GAS monoclonals Ab25 and Ab49 both led to a significant increase in Fn-binding to the bacteria while all other antibody treatments did not. To test whether this phenomenon is M protein-dependent we repeated the assay with a M protein knock-out mutant SF370ΔM (Fig. 2b). This led to no significant difference in Fn-binding. To be sure that the antibodies themselves were not able to bind Fn, we performed an ELISA with Ab coated wells. We saw that no Fn was bound to the Abs unless GAS was present (Fig. S1). To understand how polyclonal antibodies and other serum proteins can influence monoclonal induced Fn-binding we incubated SF370 in varying concentrations of pooled serum as well as pooled saliva (Fig. 2c). Monoclonals were supplemented as an additional treatment and fibronectin binding to GAS was assessed by flow cytometry. Fn-binding by GAS incubated in high concentration serum (80%) was blunted and no difference could be seen between the treatments. It is, however, noteworthy that higher serum concentrations led to higher background binding of Fn due to generally higher Fn concentrations. At lower serum concentrations (<10%) similar trends could be seen as in Figure 2a. At 1% serum and in saliva, Ab25 and Ab49 treatment consistently resulted in an increased binding of serum Fn by GAS. However, Fn-binding was only significantly increased for both Ab25 and Ab49 in saliva. In 1% serum, only Ab49 led to a significant increase in Fn-binding. We also wanted to evaluate the effect of pooled polyclonal antibodies when simultaneously applied with the anti-GAS monoclonals (Fig. S2). We therefore treated SF370 with Fn combined with IVIG (100 μg/ml) and varying concentrations of Ab25 – one of the monoclonals which consistently led to Fn-binding. As a control, bacteria were treated with only IVIG. For normalization, SF370 was left untreated, whereby only Fn was added. The signal of GAS-associated fluorescent Fn was assessed by flow cytometry. Both control samples resulted in low, background levels of GAS-bound Fn. GAS treated with Ab25 (3.75 μg/ml) led to a clear increase in bound fibronectin. When the same amount of Ab25 was used in conjunction with a much higher concentration of IVIG this led to a three-fold decrease. An increase, similar to that seen when GAS was treated with only Ab25 was observed when 7.5 μg/ml were applied with IVIG. 15 μg/ml of Ab25 with IVIG led to an increase of GAS-bound Fn compared to untreated GAS. Finally, we assessed the Fn-binding affinity to GAS with and without Ab25. We found that while Fn alone bound with a medium-high affinity of 50 nM-1, the addition of 20 μg/ml of Ab25 led to a binding affinity of KD 10 nM-1. This constitutes high-affinity binding and an around 5-fold increase in Fn-binding compared to Fn alone (Fig. 2d). However, the fit of the binding data was not as good for lower concentrations, where there seemed to be a much larger relative difference in Fn-binding, indicating that the binding difference might be underestimated. Moreover, we found that titrating the concentration of Ab25 leads to a positive modulation of Fn-binding where increasing the concentration of Ab25 leads to more Fn being bound (Fig. S3). In summary, we found that certain monoclonal antibodies targeting the M protein are able to induce increased Fn-binding to GAS while non-specific or pooled antibodies were not. When these antibodies were used in conjunction with saliva or low concentration plasma, we saw a similar phenotype which was diminished when higher concentrations of plasma were used. We saw that while pooled antibodies could dampen the increase in Fn induced by our monoclonals, this effect could be outcompeted. Finally, Fn-binding affinity measurements showed that Ab25 allows Fn to bind with increased affinity to the bacterial surface.

**Fig. 1.**
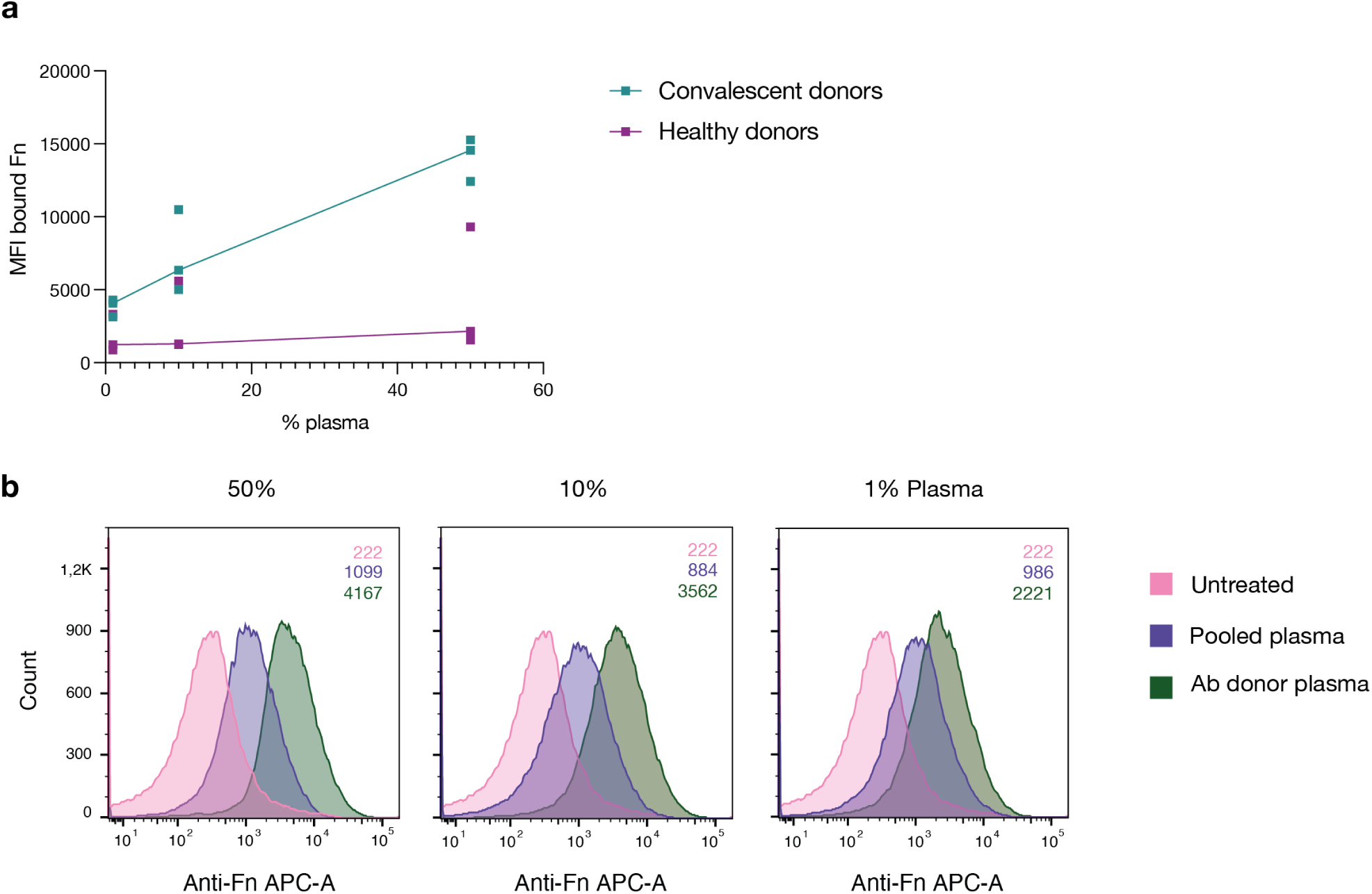
Convalescent patient plasma enhances Fn binding to GAS. **a**, GAS was incubated in plasma from patients who had recently recovered from a severe GAS infection (convalescent donors) as well as in plasma from healthy controls. This was done at three different dilutions (50, 10, and 1%). All healthy donor plasma, except one, led to greatly reduced Fn-binding to GAS compared to the convalescent donor plasma. Each point represents a distinct donor, and the line shows the respective median. **b**, GAS incubated in plasma derived from the original donor of AB25, 32, and 49 led to a dose-dependent increase in bound Fn compared to both untreated bacteria and pooled plasma (MFI of bound Fn displayed in the upper right corner). GAS was incubated in three different plasma dilutions in PBS (50, 10, and 1%). Each histogram corresponds with 20,000 bacterial events as assessed by flow cytometry. All data for this figure was acquired by flow cytometry.

**Fig. 2.**
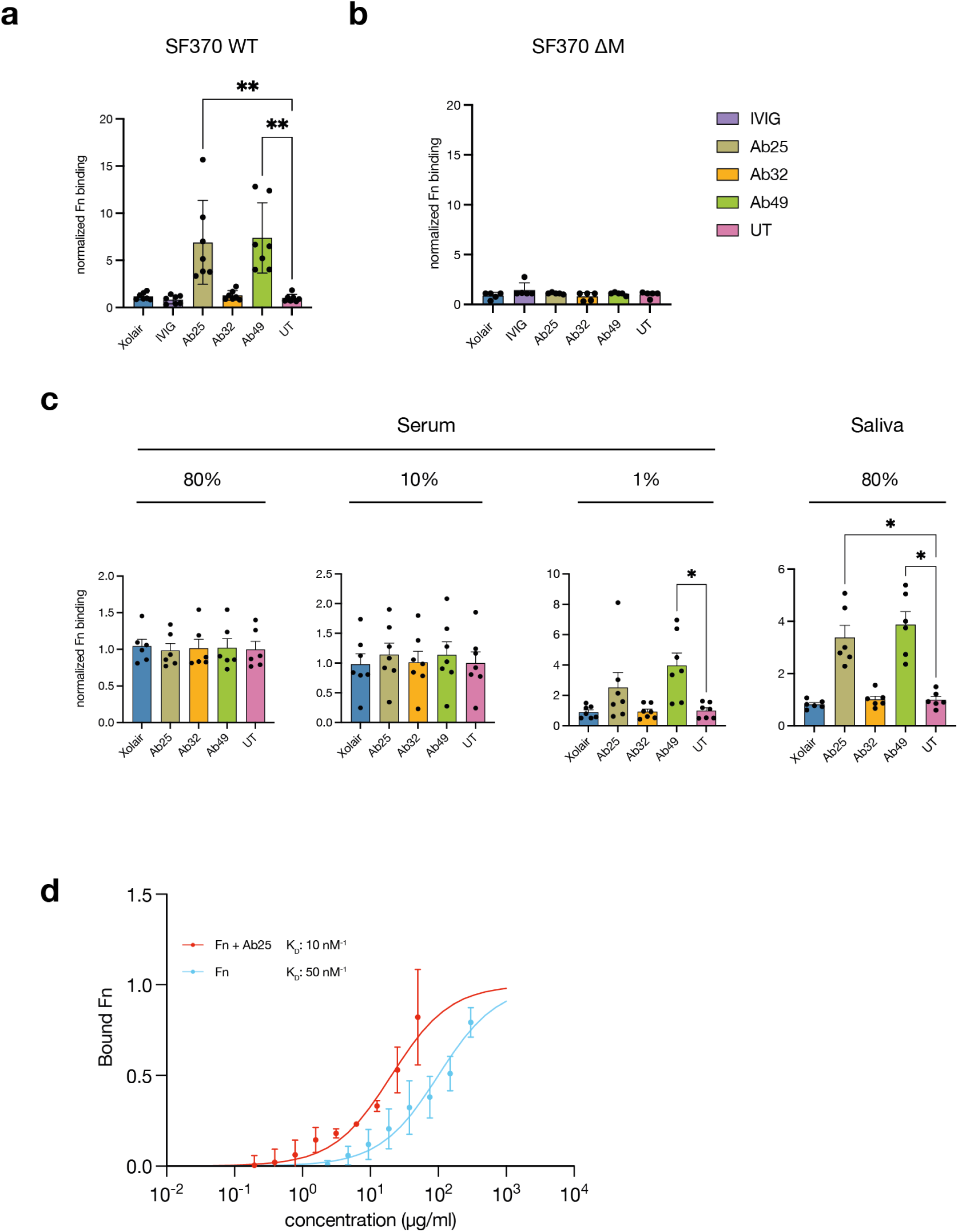
Certain monoclonal antibodies induce increased Fn-binding to M1 GAS in an M protein-dependent manner. **a**, the M-specific monoclonals Ab25 and 49 lead to a significant increase in Fn-binding to SF370 GAS compared to the untreated control (UT, no additional antibody treatment). The non-binding control (Xolair), pooled intravenous antibodies (IVIG), the M-specific monoclonal Ab32 did not. **b**, the M1 protein knock-out mutant SF370 ΔM was treated with Fn alone (UT) or in combination with Xolair, IVIG, and the M-specific monoclonals (Ab25, 32, and 49). All antibodies and Fn were added at 20μg/ml. No significant difference in Fn-binding was observed for any of the treatments compared to the untreated control. **c**, lower Ab background levels (80% saliva and 1% serum) result in an increased GAS Fn-binding compared to the untreated control (UT, no additional antibody treatment) when Ab25 and 49 are substituted while this was not seen with other Ab treatments. Higher antibody titers found in 80% and 10% serum reduced the effect substituted Abs had on GAS Fn-binding (all Abs were added at 20 μg/ml).**d**, Fibronectin binding affinity increases around 5-fold in the presence of Ab25. Binding curves of fibronectin with and without 20 ug/ml Ab25 give an estimate of affinity to protein M1. Fibronectin binding was measured using flow cytometry. The figure shows measured binding with fitted ideal binding curves as a function of the total fibronectin concentration. N=3 for all concentration points. Dissociation constants (KD) for the curves are given in the plot, together with a confidence interval calculated using the Bootstrap method. Error bars represent the SEM. Statistical significance was assessed using Kruskal-Wallis combined with Dunn’s multiple comparisons test and * denotes p 0.05 and ** for p 0.005.

### Fn binds to the upper domain in M proteins and binding co-varies with specific antibodies

While it was clear that certain antibodies are necessary to induce high-affinity Fnbinding to GAS we had not yet assessed what this means for the Fn-binding binding capacity of individual M proteins. To assess this, we employed super-resolution fluorescence colocalization microscopy (Fig. 3a). Colocalization was assessed between Fn and a corresponding antibody treatment (monoclonals Ab25 and Ab49, or polyclonal antibodies isolated from convalescent patient plasma (CAbs)). Images of single bacterial cells from each treatment group were acquired using SIM microscopy, and protein binding to the bacteria was individually assessed. We found that Ab binding uniformity varied greatly depending on the respective Ab treatment and the bacterial cell being assessed. However, we found that, regardless of Ab treatment, the overwhelming majority of assessed bacteria showed Fn signal that clearly colocalized with antibodies bound to the bacterial surface. Additional representative images from the dataset of assessed bacteria can be found in the supplementary (Fig. S4). Furthermore, we wanted to assess if the Fn-binding site on the M1 protein is truly in the N-terminal A-B repeats as suggested previously (21) and whether this binding site remains the same if Fn-binding is induced by bound Abs. To assess this, we used a microscopy-based site determination method (28). This allowed us to measure the distance of a target protein from the cell wall with nanometer-scale precision (Fig. b-c). Distance measurements done with control proteins, that bind to known sites on M1, served as landmarks and allowed us to extrapolate which relative region fibronectin was binding to. First, we assessed the Fn-binding induced by isolated convalescent plasma antibodies (Cabs) and low-affinity Fn-binding by treating with a high Fn concentration (500 μg/ml). As a control, we assessed the binding distance of fibrinogen (Fb) (Fig. 3b). The measurements showed that both CAb induced, high-affinity Fn-binding and low-affinity binding of highly concentrated Fn led to binding in the same region of M (median binding distance: 26.5 and 22.4 nm respectively). This lay below the binding distance of Fb (median binding distance 46.4 nm) which is known to have multiple binding domains in both the A and B repeats and therefore also displays a larger data spread (29). Next, we measured the binding distance for Fn bound due to Ab25 or Ab49 and as controls assessed the binding distance of the antibodies themselves which have both been shown to bind the central S region (27) (Fig. 3c). We found that again, Fn was being bound in a similar location to the previous Fn-binding distance measurements (median binding distance: 35.2 and 27.7 nm). The corresponding Ab binding measurements presented smaller distance values (median binding distance: 11.9 and 13.5 nm), verifying that Fn was binding above both Ab binding epitopes on M – the constant region of M1. In summary, colocalization scoring was done with the two functional monoclonals as well as polyclonal antibodies purified from convalescent patient plasma. It showed that antibody binding to the bacterial surface was highly heterogeneous and that most bacterial cells assessed showed clear colocalization between areas with bound antibodies and Fn. Site determination analysis found that, regardless of how the binding is realized, Fn will bind to a region on the M protein which lies between the central S region and the Fb binding region in the A repeats.

**Fig. 3.**
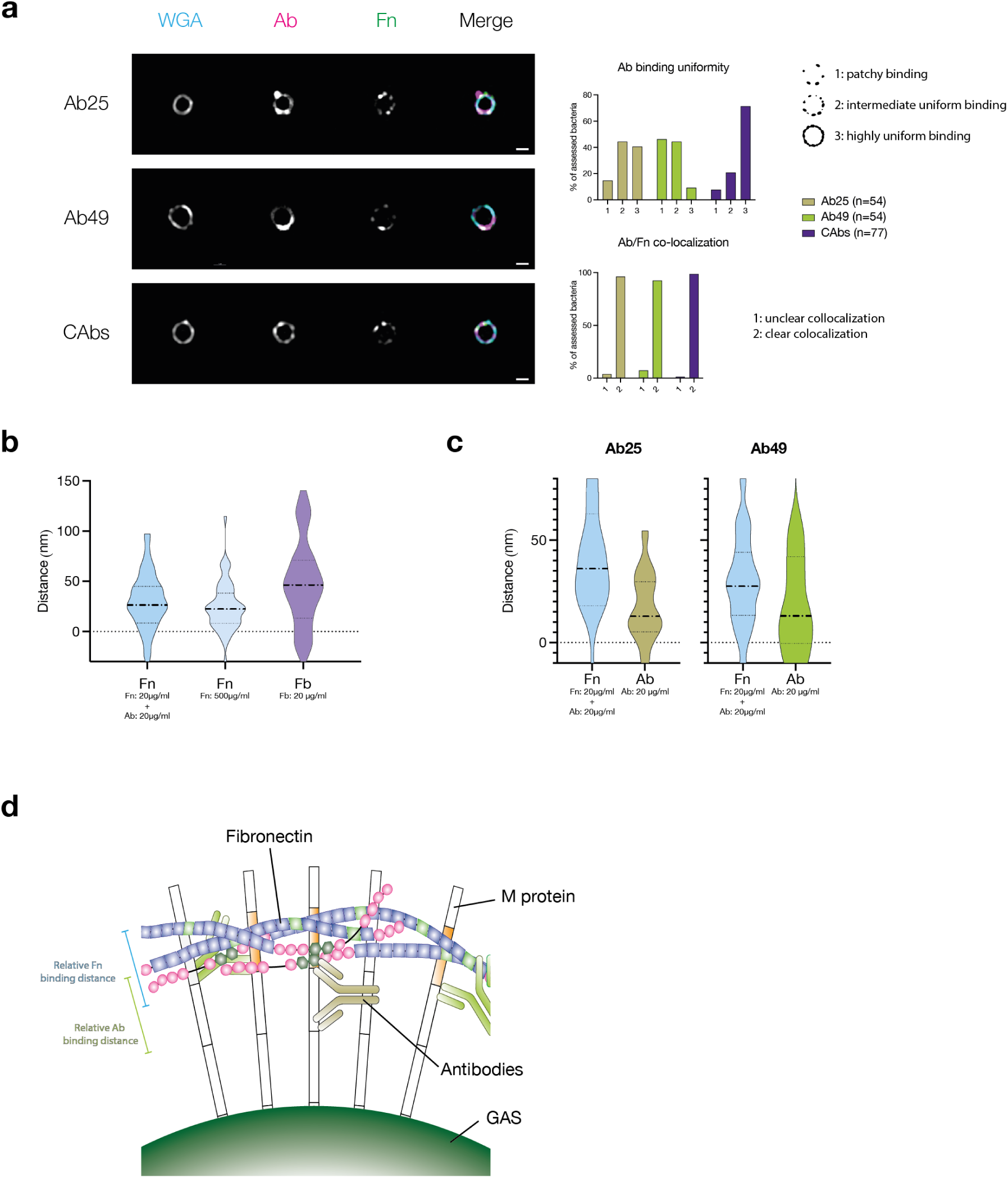
Fn binds to the upper domain in M proteins and binding co-varies with specific antibodies. **a**, Representative fluorescence colocalization images of SF370 imaged with a N-SIM microscope. The cell wall was stained wheat germ agglutinin (WGA). The bacteria were then treated with Fn and one of 3 antibody treatments. These were Ab25 (top), Ab49 (middle), and antibodies derived from a GAS infection convalescent donor (CAbs, bottom). On the right side, merged images of the WGA-stained cell wall, Abs, and Fn are represented by cyan, magenta, and green respectively. The scale bar represents 1 μm. To the right of the image panel, the scoring data from the assessed bacterial cells is displayed. At the top, the antibody binding uniformity scoring is displayed, and below that the colocalization scoring results are shown. The score value attribution can be seen to the right of the graphs. **b**, Fn and fibrinogen (Fb) binding site distance measurements from the bacterial surface are displayed as violin plots. Fn-binding resulting from CAb treatment and low-affinity Fn-binding, achieved by incubating the bacteria in high concentrations of Fn are displayed next to the results from a binding distance control Fb. Both Fn-binding measurements showed that Fn consistently bound below Fb on the M protein. **c**, Violin plots showing binding site distance measurements of Fn and corresponding Ab used to induce said Fn-binding (Ab25, Ab49). Regardless of the antibody treatment, the measured Fn-binding distance was very similar and was in both cases larger than the Ab binding distance. **d**, Representative illustration showing a proposed binding model of Abs inducing Fn-binding to surface-bound M protein. The scale bars represent relative binding distances attained from binding distance measurements.

### Monoclonal antibodies induce enhanced Fn-binding in several tested M types

We had now confirmed that Fn was being bound in the N-terminal variable region of M and that certain antibodies, binding to the central conserved region, could induce high-affinity Fn-binding. We knew that, due to their binding site, our antibodies broadly react with multiple M types with differing potency (27). This presented us with the opportunity to assess how generalizable the described phenomenon is amongst various M strains and if we would see a similar pattern than we had previously seen with antibody binding to GAS. We tested our monoclonal antibodies’ ability to induce Fn-binding with a total of four additional GAS strains expressing various M types (M5, M12, M79, M89) (Fig 4a). We found that compared to the M1 strain, all four other tested strains exhibited far higher background levels of Fn-binding between 50 and 150 times more Fn was bound by untreated (UT = no Ab treatment) GAS strains compared to the M1 strain. The strains M1, M79, and M89 all bound substantially higher amounts of Fn when exposed to our monoclonal antibodies (Ab25 and Ab49) while the M5 and M12 strains did not. This matched the pattern seen for antibody binding to the strains (27). The Fn-binding data were additionally normalized to the UT controls for each strain which allowed for better visualization of the effectiveness of each Ab treatment on a separate strain. We found that, in terms of fold change, the M1 strain was impacted most by the Ab treatment where, on average, Ab25 led to 40-fold more Fn being bound. While we saw a significant increase in Fn-binding for the strains M79 and M89, the high background levels of bound Fn in the UT group made the effect less pronounced. Next, we wanted to better understand how these distinct M types could be binding Fn under the influence of our bound monoclonals. We, therefore, decided to sequence the various M proteins expressed by these strains. Since we had previously determined that Ab-induced Fn-binding occurs in the variable A/B repeats, we aligned the amino acid sequences of all five strains (Fig. 4b). To allow for better alignment, we included the signal peptide in the analysis. Our alignment showed that while there were naturally strong homologies within the signal peptide sequence, the A/B repeats showed almost none at all. The binding epitopes of both Ab25 and Ab49 have been framed in red and green. For better comprehension, we have included a cartoon of M protein to illustrate the Ab binding sites and proposed relative Fn-binding site. (Fig. 4c) We used a percent identity matrix to compare sequence similarities between M types (Fig. 4d). It shows that even though M1, M79, and M89 all bound increased levels of Fn under the influence of our monoclonals, their variable A/B were not substantially more similar than M5 and M12, which did not. An only slightly higher semblance was observed between M79 and M89. Fn-binding data comparing 5 different M types showed that different M strains bind vastly different base levels of Fn and that our monoclonals targeting the conserved region of M led to significantly increased binding in 3 of 5 strains. This matched the capacity of the monoclonals to bind to the strains themselves as seen in a previous study (27). By comparing the amino acid sequences of the variable A/B regions of the various M proteins, we found that the ability to bind Fn to the M protein under the influence of bound antibodies did not correlate with substantial sequence homology.

**Fig. 4.**
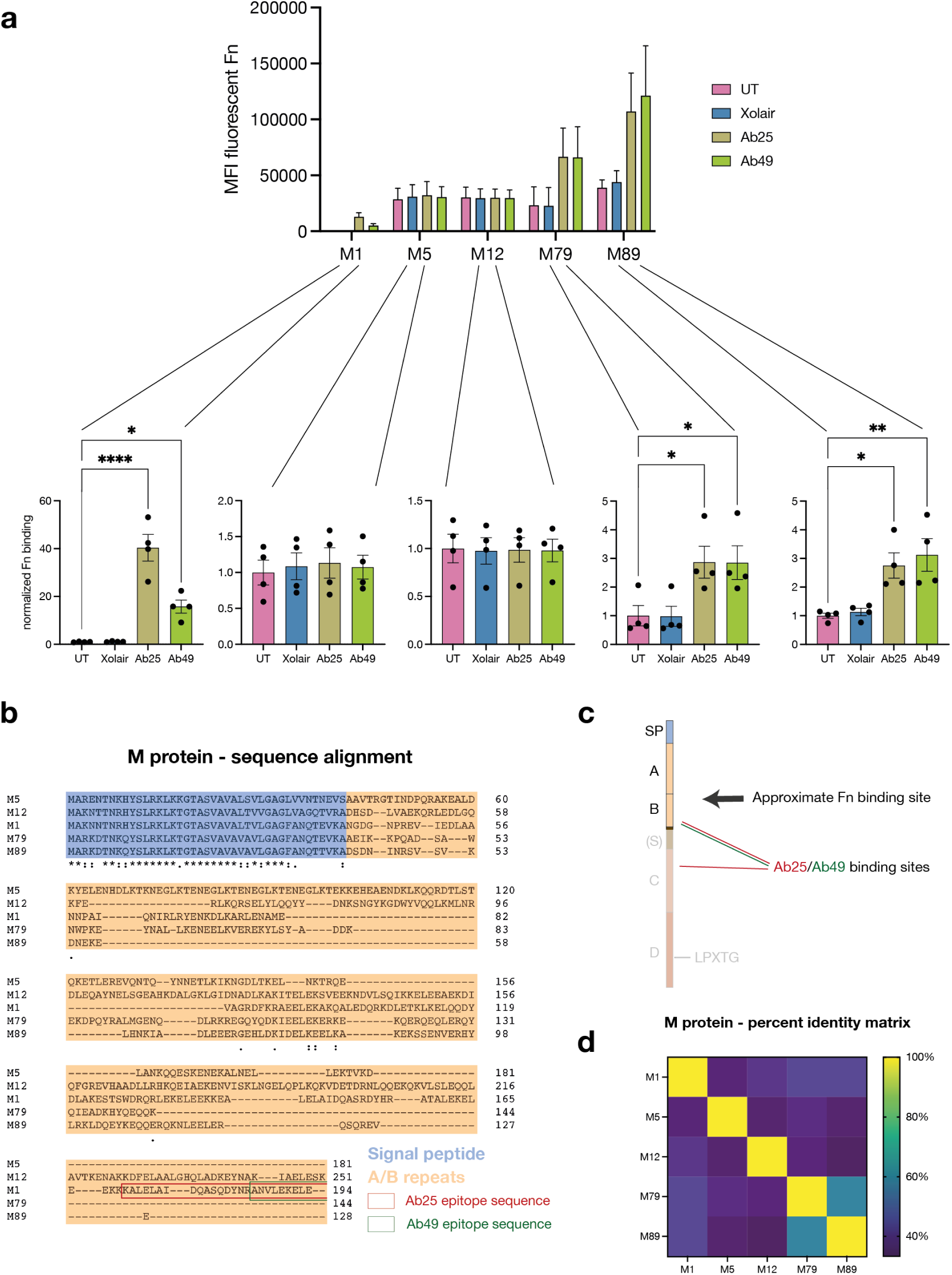
Monoclonal antibodies induce enhanced Fn-binding in several tested M types. **a**, Five GAS strains expressing distinct M types (M1, M5, M12, M79, and M89) were exposed to Fn in combination with Ab treatments. All strains showed variable levels of background Fn-binding when no treatment (UT) or no specific antibody was present. For 3 of the 5 tested strains (M1, M79, and M89), treatment with the monoclonals Ab25 or 49 resulted in a significant increase in Fn-binding compared to the untreated control. All antibodies were added at 20μg/ml. The data for this graph was acquired by flow cytometry. **b**, Amino acid sequence alignment of the various tested M types. The sequences include the signal peptide (blue), A/B regions (orange), and the binding epitope sequences of Ab25 and 49. **c**, Illustration depicting the GAS M protein, divided into its various regions whereby not all M types feature an S region. The binding sites of Ab25 and 49 are shown in red and green respectively, whereby Ab25 binds to the M protein with both of its Fab domains in dual-Fab cis binding conformation (27). Site localization analysis (Fig. 3b-c) revealed that Fn binds above both antibodies. **d**, Percent identity matrix showing the similarity of the aligned amino acid sequence of all tested M types. The data is shown as a heatmap. Error bars represent the SEM. Statistical significance was assessed using one-way ANOVA combined with Dunnett’s multiple comparisons test and * denotes p 0.05, ** for p 0.005, and *** for p 0.001, and **** for p 0.0001.

## Discussion

It is well known that GAS possesses a multitude of Fnbinding proteins which it uses to recruit Fn to its surface – assisting in the process of colonization and infection of a host. We were surprised to find that a GAS strain of the most common M type (M1) bound only insignificant amounts of Fn when incubated in human plasma. Further, we found that incubation in plasma of convalescent patients led to a substantial increase in bound Fn instead of an expected reduction due to antibody interference. Using purified Fn and antiM monoclonal antibodies we found that it was in fact certain M protein targeting antibodies that led to this increase in Fn-binding and that pooled antibodies from healthy donors (IVIG) led to no such increase. When GAS was treated with varying concentrations of pooled serum in combination with the anti-M monoclonals, we saw that higher serum concentrations led to a dampened effect of the monoclonals on Fn-binding. This was most likely due to anti-GAS antibodies found in the serum which at high concentrations can outcompete the monoclonals. However, at lower serum concentrations, as well as in saliva, the anti-M had a pronounced effect on Fn-binding indicating that this phenomenon is a niche-specific effect. This would make sense since GAS most commonly infects and colonizes humans in the pharynx – an environment with low concentrations of IgG. This niche specificity, also reported for Fc binding function in GAS (30), could also explain why we see slight differences between 1% serum and saliva. While these most likely contain comparable antibody concentrations, there are many serum proteins, not found in saliva, which could be interfering in unknown ways. While the pooled antibodies (IVIG) bound to GAS, we found that far lower concentrations of our anti-M monoclonals could outcompete this binding indicating that a small antibody subpopulation is sufficient to induce this phenomenon. In fact, we found that one of our three anti-M monoclonals, Ab32, did not lead to such an effect. This could be due to this antibody’s binding epitope – since Ab25 and 49 essentially share a binding epitope. It is possible that this phenomenon only occurs when antibodies bind to a very specific region of the M protein. Affinity measurements revealed that Fn alone bound to GAS with medium-high affinity. However, this binding affinity could be increased 5-fold with the addition of Ab25.

To show that antibodies had to be bound to M protein in order for efficient Fn-binding to occur, we utilized fluorescence colocalization microscopy. Even in cases of non-uniform surface binding of the antibodies, we found that Fn overwhelmingly colocalized with bound antibodies. This indicates that a direct interaction between M, antibodies, and Fn is necessary to trigger the observed effect. Next, we wanted to assess whether, in the presence of antibodies, Fn still bound to the N-terminal region of M as previously described (21). We employed a microscopy-based site localization method to assess how far from the bacterial surface Fn was binding. We found that regardless of the antibody treatment used to trigger it, or if it was only Fn-binding by itself, Fn-binding always occurred above our reference antibodies which bind to the conserved S region of M1. Finally, since we now knew that the Fn-binding was occurring in the variable N-terminal region of M we wanted to assess if our monoclonals targeting a conserved region would induce the same response in other M types. We found that, of the four additionally tested M types, two bound more Fn in the presence of the monoclonals. Amino acid sequence analysis revealed that the variable A-B repeats of all five strains differed significantly showing no clear sequence homologies. This indicates that, as previously shown for the functionality of other bacterial surface proteins, through convergent evolution, sequence homology is not a prerequisite for identical function (31).

*Streptococcus pyogenes* was one of the first bacterial pathogens proven to bind the host protein Fn to its surface. It quickly became clear that this capability must be key to stable colonization of a host and that it is of immense benefit to the bacterium (22). Upon more in-depth study highly complex immune-subversive mechanisms between GAS-bound Fn and host cells came into focus (11, 32, 33). In recent years many responsible Fn-binding proteins have been discovered and in fact it is often the case that GAS strains express multiple heterologous Fn-binding proteins simultaneously (4). Protein production is a costly process in terms of energy consumption and so, from an evolutionary standpoint, it makes no sense for bacteria to express proteins that do not benefit them. This makes it clear that Fn-binding must have been a crucial function during the millennia of co-evolution with the human immune system. We found that SF370, the M1 strain used in this study, was only able to bind Fn very weakly. This, despite the fact that it possesses the Fn-binding protein FbaA (34) and has been shown to be able to bind Fn to its M protein (21, 32). We must therefore ask ourselves: what is the benefit of these Fn-binding proteins if they only bind Fn so weakly? The findings detailed in this study indicate that certain M1 GAS have evolved to ensnare and benefit from antibodies proteins built by the human immune system to be inherently damaging to them. It is possible that M1 strains are partly so successful since this mechanism gives them an evolutionary advantage? It would be interesting to further assess how this affects the virulence of such strains. Previous work found that complementing an M1-type strain with the Fn-binding protein F1 resulted in attenuated virulence invivo (35). At first, this may seem contradictory since GAS-mediated Fn-binding has so clearly been linked to virulence (11, 20, 33, 36). However, this may be an indication of the importance of Ab enhanced M protein-mediated Fn-binding. If certain M1 strains have adapted to be highly reliant on a more energy-efficient means of binding Fn, such as binding Fn with the assistance of binding antibodies, then the forced expression of a superfluous protein could lead to metabolic dysregulation resulting in lower fitness and thus virulence. This could also be the case for the investigated M1 type strain SF370. We found that it was very inefficient at binding Fn even though it processes the gene for FbaA. Could evolutionary reliance on Ab-mediated Fn-binding to the M1 protein have led to a low expression of this specialized Fn-binding protein? Antibodies inducing such an effect seem to target the conserved central region of the M protein. Intriguingly it is this region that has been shown to attract the strongest humoral immune responses (14, 15). Due to the conserved nature of this region, antibodies targeting it will be reactive towards many M types. We have seen that other M strains such as M79 and M89 bound more Fn in the presence of the antibodies. However, it is more difficult to understand the phenomenon’s usefulness for these strains since they already express highly efficient Fn-binding proteins. It, however, explains why the Fn-binding potential of these two M proteins seems to have been overlooked up until now.

While the observed effect of these antibodies on M protein-mediated Fn-binding is enticing, the exact mechanism remains unknown. M protein is highly structurally complex and its specific conformation – dictated by environmental factors such as temperature and pH alterations – has been shown to have substantial effects on its function (34,35). It has previously been shown that certain antibodies, upon binding, can lead to conformational shifts in the target epitope. Such findings include reports of antibodies, upon binding, leading to the inactivation or re-activation of enzymes, allowing synergistic binding between antibodies or altering the structure of a ligand to enhance binding to its receptor (36–39). It is therefore plausible that the binding of an antibody to a neighboring domain on M is leading to an allosteric shift which then allows for higher affinity Fn-binding. Another explanation could be that the binding antibody is locking the bound Fn in place by steric hindrance. Further studies will be needed to better understand the precise mechanisms which allow the observed enhanced binding to occur. Finally, it will be critical to study the physiological implications caused by this phenomenon. GAS has already been linked to many post-infection autoimmune sequelae such as rheumatic carditis, glomerulonephritis, and Sydenham chorea. All these diseases are mediated by antibodies which are thought to be generated due to molecular mimicry between the M protein constant regions and various human tissues (40). As previously mentioned, the M protein C repeats have been seen to act as an immunogenic flag, actively drawing in an antibody response (14, 15). It has not fully been understood why GAS would benefit from actively encouraging the generation of such antibodies since some can be detrimental to the bacteria (27). However, our findings show that it is exactly these antibodies which may allow the M protein to efficiently bind Fn, directly benefiting the bacteria. Moreover, it would be important to understand how it is beneficial for the bacteria to possess a protein with a function which can be activated through outside influence. It is possible that this phenomenon acts as a sensing mechanism – triggering accumulation of Fn on the bacterial surface just when the immune system begins mounting an immune response. This could potentially allow the bacteria to ‘react’ to a forthcoming hostile immune response. This would not be the first report of GAS exhibiting mechanisms to ‘sense’ environmental processes. Previous observations found that by employing the major GAS surface protein SclA/Scl1, GAS can adapt and respond to the host’s wound environment by selectively binding wound associated isotypes of Fn (44). Future studies exploring the physiological impacts of antibody mediated binding of Fn to M protein could prove essential in better understanding the highly complex relationship between GAS and the immune system.

## Materials and Methods

### Bacterial strains, growth, and transformation

Five different GAS strains, expressing distinct M types were used for this study (M1, M5, M12, M79, M89). Both M1 and M5 expressing strains, SF370 and Manfredo respectively, have been thoroughly studied in previous studies (34, 45) The other strains (M12, M79, M89) were clinical isolates derived from patients with severe GAS infections. These specific Mtype strains were chosen since they were previously used to test the cross-strain reactivity of the monoclonals. In this assay, M5 and M12 showed the weakest reactivity while M79 and M89 showed the strongest. (27) The GAS strains were grown statically in Todd-Hewitt Yeast media (THY) at 37°C, 5% CO2. The strains were maintained on THY agar plates for three weeks before being replaced with a plate freshly streaked from −80°C stocks. For binding experiments, bacteria were grown to logarithmic phase by diluting an overnight culture 1:20 in fresh THY. After dilution bacteria were grown until they were in the mid-log phase – around 2 to 2.5 hours or until they reached an OD600 of 0.4. For the transformation of GFP expressing SF370 and the M1 knock-out mutant strain M were washed in ice-cold water to make them electrocompetent. These bacteria were electroporated with 20 μg of the pGFP1 plasmid and after a 1-hour recovery period, the bacteria were plated on THY plates supplemented with erythromycin. Successful transformants were tested for fluorescence under ultraviolet light. Heat-inactivation of the bacteria was done by growing them to mid-log as described previously. Thereafter, they were washed in PBS before being transferred to ice for 15 minutes. Next, the bacteria were heat-shocked at 80°C for 5 minutes and immediately transferred back to ice for another 15 minutes. For site localization analysis SF370 GAS was grown to mid-log as previously described. The culture was then centrifuged and washed in 10 ml of PBS. The bacteria were then resuspended in 1ml of PBS. The bacterial cell wall was, depending on the experiment, either stained with AlexaFluor (AF) 488 or AF594 conjugated WGA. After staining the bacteria were sonicated and then fixed in 1% PFA for 30 minutes at RT. After fixing, Tris was added for a final concentration of 333 mM to quench PFA before centrifugation. This reduced bacterial clumping during centrifugation.

### Plasma and serum samples

The convalescent patient samples used in Fig. 1a were received from colleagues working at the infectious disease department. We were not given information beyond the fact that all three patients had suffered and recovered from severe invasive GAS infections. The healthy donor plasma for this panel was derived from healthy donors who had no symptoms at the time of venous puncture. For Fig. 1b plasma derived from the GAS convalescent donor whose b-cells were originally isolated to attain the monoclonals tested in this study (Ab25, 32, 49). The pooled plasma was commercially available pooled control plasma (Affinity Biologicals Inc.). For the experiments depicted in Fig. 2c the pooled serum was purchased from Sigma. The pooled saliva was made by pooling saliva from 10 healthy donors. The saliva was then centrifuged to remove coarse debris and then filtered with a 0.2 μm syringe filter.

### Monoclonal antibodies

The monoclonal antibodies employed in this study were previously generated and investigated at length (27). All antibodies were of the subclass IgG1. Ab25, Ab32, and Ab49 were found to bind to GAS M protein with high specificity and affinity. The antibodies were also shown cross-react with M types. Cross-linking combined with mass spectrometry revealed that the binding epitopes of all three monoclonals lay close to the central S region of the M1 protein. Ab25 presented a special binding conformation whereby it bound the M protein in cis dual-Fab conformation. This means that it bound M with both its Fab domains simultaneously with epitopes on both sides of the S domain.

### Antibody purification, digestion, and assessment

Monoclonal antibody generation and purification was done as previously described (27). Antibody purification from plasma was done by incubating plasma with protein G Sepharose beads (Cytiva). The beads were incubated with the plasma for 2 hours before being transferred to a chromatography column. The beads were then washed 4 times with 10 ml of PBS without allowing the beads to dry out between or after washes. The antibodies were then eluted with 0.1 M glycine which was immediately buffered with Tris. Centrifugal filter columns (Millipore, Merk) were used to exchange the buffer to PBS.

### Fibronectin and antibody conjugation

Fn purified from human plasma (sigma) or monoclonal antibodies were fluorescently conjugated using AlexaFluor 488 and 647 Ester (Invitrogen). The conjugation and purification were done according to the manufacturer’s guidelines. Protein concentration after purification, as well as the degree of labeling, was assessed using a DeNovix DS-11 FX system.

### Fn-binding assays and flow cytometry

GFP expressing SF370 WT and ΔM, as well as M5, M12, M79, M89 GAS were grown to mid-log as previously described. The bacteria were centrifuged and thoroughly resuspended in 1 ml of PBS. For patient plasma incubation experiments (Figure 1a-b) varying dilutions of plasma were prepared in a 96-well plate whereupon 10 μl of the mid-log culture concentrate was directly added. As pooled plasma, we used frozen normal control plasma (VisuCon-F, Affinity Biologicals) which, according to the manufacturer, was collected from at least 20 healthy donors. Bacteria were incubated with the plasma for 30 min at 37°C. After incubation, the plate was centrifuged, and the bacteria were washed twice in PBS. The bacteria were stained with an anti-Fn antibody (Thermo Fisher) and incubated for 30 min at 37°C. The bacteria were then washed once to remove the primary antibody and then stained with an anti-mouse fluorescent secondary conjugated with AlexaFluor 647 (Thermo Fisher). Fibronectin binding was assessed by flow cytometry (CytoFlex, Beckman Coulter). GFP-expressing bacteria were assessed by gating for FITC positive events. This gate was then assessed for Fn median fluorescence intensity (MFI) in the APC channel. Since it was not possible to transform all M strains for the cross-strain binding experiment the bacteria were gated using SSC and FSC. The accuracy of this gating strategy was confirmed by comparing both gating methods with GFP-expressing strains. The pooled serum and saliva experiment (Figure 2c) was done following a similar protocol. Pooled human serum (Sigma) dilutions and pooled saliva were prepared in a 96-well plate. The anti-GAS monoclonals Ab25, Ab32, and Ab49 as well as the non-binding control Xolair (Novartis) was added at 20 μg/ml before 10μl of the concentrated mid-log SF370 culture was added. As an untreated control (UT), PBS was added instead of monoclonal antibody treatment. The rest of the experiment was done following the same protocol as mentioned previously. After the acquisition, the MFIs were normalized to the results attained from the untreated samples. The Fn binding experiments only employing purified antibodies (Fig. 2a-c and Fig. 4a) were done by preparing a plate with the corresponding antibodies (20 μg/ml) and AlexaFluor 647-Fn (20 μg/ml). As an untreated control (UT), only PBS was used. 80 μl of a non-concentrated mid-log culture of the corresponding bacteria was added to each well. The bacteria were incubated for 30 minutes at 37°C. After incubation, the bacteria were washed twice with PBS before assessment of bound Fn by flow cytometry as detailed above.

### Affinity measurements

10 ml of a SF370 culture were concentrated into 1000 μl of PBS, and 10 microlitres of bacteria were added to three 1.5ml tubes – one for each binding curve. 300 μg/ml of AlexaFluor 647-conjugated fibronectin was added to the first tube. 50 μg/ml of AlexaFluor 647conjugated fibronectin and 20 μg/ml of Ab25 were added to the second tube. 1 μg/ml of AlexaFluor 647-conjugated fibronectin and 20 μg/ml of Ab25 were added to the third tube. All tubes were incubated on shake at 4 degrees Celsius for 30 mins. Bacteria and all constant protein concentrations for each binding curve were added to wells of a 96-well plate – 20 μg/ml of Ab25 in each well for the second binding curve and 20 μg/ml fibronectin in each well for the third binding curve. Serial dilutions were made with the prepared wells using the tube samples as maximum concentration points. The well plate was incubated on shake for 30 mins at 4 degrees Celsius. Flow cytometric acquisition was performed using a CytoFLEX. Theoretical fit was performed in MATLAB using a weighted least squares method for an ideal binding curve with the dissociation constant as an unknown variable. The accuracy of predicted affinity estimates was calculated using the Bootstrap method (46) and is the confidence interval calculated from 50 resamplings of the acquired data.

### SIM imaging

For the fluorescent colocalization microscopy experiments as well as determination of the Fn-binding site, the bacteria were prepared as mentioned above. For the colocalization experiments, the cell wall of SF370 GAS was stained using AlexaFluor 594-conjugated wheat germ agglutinin (WGA). The Fn was conjugated with AlexaFluor 488 and the bound antibodies were visualized using a Fab antihuman IgG Fab secondary conjugated with AlexaFluor 647 (Jackson ImmunoResearch). The site localization experiments were done with SF370, whereby the cell wall was stained with AlexaFluor 488-conjugated wheat germ agglutinin (WGA). They were then opsonized with AlexaFluor 647-conjugated Fn and corresponding non fluorescently labeled antibodies. All treatments were used at 20 μg/ml except high Fn, used to show low-affinity Fn-binding, for which 500 μg/ml of Fn was used. To ensure thorough opsonization the bacteria were incubated with their corresponding treatments for 30 minutes at 37°C while shaking. After incubation, the bacteria were washed once in PBS. Samples were mounted on glass slides using Prolong Gold Antifade Mountant (Invitrogen) with 1.5H coverslips. Single bacteria were manually identified and for site localization determination they were imaged with a time series of 15 images per channel. Images of single bacteria were acquired using a Nikon N-SIM microscope equipped with a LU-NV laser unit, CFI SR HP Apochromat TIRF 100X Oil objective (N.A. 1.49), and an additional 1.5x magnification. The camera used was ORCAFlash 4.0 sCMOS camera (Hamamatsu Photonics K.K.). Reconstruction was done with Nikon’s proprietary SIM software included in NIS Elements Ar (NIS-A 6D and N-SIM Analysis). The analysis pipeline for site determination was implemented in Julia and is available on GitHub (28). Representative illustrations showing hypothetical protein binding to M were made in Adobe Illustrator.

### M strain sequencing and amino acid alignment

The genomic DNA of the 5 assessed M strains were isolated and purified using the Wizard Genomic DNA Purification Kit (Promega). The growing of strains, bacterial lysis, and DNA were done according to the manufacturer’s guidelines. After isolation, the DNA was sequenced using the TruSeq DNA PCR-Free platform from Illumina. The M protein amino acid sequences were aligned and analyzed using Clustal Omega (1.2.4).

### Graphs and statistical analysis

All graphs and statistical analysis were done in GraphPad Prism (9.3.1.). To assess statistical differences in the Fn-binding to SF370 GAS induced by various anti-M monoclonals (Fig. 2a-c) we employed the Kruskal-Wallis H test combined with the Dunn multiple comparisons post hoc test. For the assessment of statistical differences in Fn-binding to various M-type GAS strains (Fig. 4a) we employed an ANOVA combined with Dunnett’s post hoc multiple comparisons test.

## Acknowledgements

SW, VKA, OA, and WB are funded by the Royal Physiographic Society. PN is funded by the Knut and Alice Wallenberg Foundation (KAW) and the Swedish Research Council (VR). We thank Anna Bläckberg and Magnus Rasmussen for giving us access to GAS convalescent patient plasma. We thank Oonagh Shannon for giving us access to the GAS clinical isolate strains. We thank the Lund University Bioimaging Centre (LBIC) for use of fluorescence microscopes. We thank the center for translational genomics (CTG) for assistance sequencing the various GAS strains used in this work. We thank Berit Olofsson, Arman Izadi, and Johannes Kumra Ahnlide for technical assistance.

## Author contributions

Conceptualization: SW, WB, and PN. Experimentation and data analysis: SW, VKA, and OA. Writing original draft: SW and PN. All authors contributed to reading and editing the final manuscript.

## References

1. Carapetis JR, Steer AC, Mulholland EK, Weber M. 2005. The global burden of group A streptococcal diseases. The Lancet Infectious Diseases 5:685–694.

2. Walker MJ, Barnett TC, McArthur JD, Cole JN, Gillen CM, Henningham A, Sriprakash KS, Sanderson-Smith ML, Nizet V. 2014. Disease manifestations and pathogenic mechanisms of group A Streptococcus. Clinical Microbiology Reviews 27:264–301.

3. Labat-Robert J. 2012. Cell-Matrix interactions, the role of fibronectin and integrins. A survey. Pathologie Biologie 60:15–19.

4. Hymes JP, Klaenhammer TR. 2016. Stuck in the middle: Fibronectin-binding proteins in gram-positive bacteria. Frontiers in Microbiology https://doi.org/10.3389/fmicb.2016.01504.

5. Pankov R. 2002. Fibronectin at a glance. Journal of Cell Science 115:3861–3863.

6. Preissner KT, Chhatwal GS. 2014. Extracellular Matrix Interactions with Gram-Positive Pathogens. Gram-Positive Pathogens, Second Edition 89–99.

7. Brouwer S, Barnett TC, Rivera-Hernandez T, Rohde M, Walker MJ. 2016. Streptococcus pyogenes adhesion and colonization. FEBS Letters. Wiley Blackwell https://doi.org/10.1002/1873-3468.12254.

8. Wang B, Yurecko RS, Dedhar S, Cleary PP. 2006. Integrin-linked kinase is an essential link between integrins and uptake of bacterial pathogens by epithelial cells. Cellular Microbiology 8:257–266.

9. Cue D, Southern SO, Southern PJ, Prabhakar J, Lorelli W, Smallheer JM, Mousa SA, Patrick Cleary P. 2000. A nonpeptide integrin antagonist can inhibit epithelial cell ingestion of Streptococcus pyogenes by blocking formation of integrin α5β1-fibronectin-M1 protein complexes. Proc Natl Acad Sci U S A 97:2858–2863.

10. Hauck CR, Borisova M, Muenzner P. 2012. Exploitation of integrin function by pathogenic microbes. Current Opinion in Cell Biology 24:637–644.

11. Yamaguchi M, Terao Y, Kawabata S. 2013. Pleiotropic virulence factor - Streptococcus pyogenes fibronectin-binding proteins. Cellular Microbiology 15:503–511.

12. Li Y, Rivers J, Mathis S, Li Z, Velusamy S, Nanduri SA, Van Beneden CA, Snippes-Vagnone P, Lynfield R, McGee L, Chochua S, Metcalf BJ, Beall B. 2020. Genomic Surveillance of Streptococcus pyogenes Strains Causing Invasive Disease, United States, 2016–2017. Frontiers in Microbiology 11:1–13.

13. Frost HR, Davies MR, Velusamy S, Delforge V, Erhart A, Darboe S, Steer A, Walker MJ, Beall B, Botteaux A, Smeesters PR. 2020. Updated emm-typing protocol for Streptococcus pyogenes. Clinical Microbiology and Infection 26:946.e5-946.e8.

14. Lannergård J, Gustafsson MCU, Waldemarsson J, Norrby-Teglund A, Stlhammar-Carlemalm M, Lindahl G. 2011. The hypervariable region of streptococcus pyogenes M protein escapes antibody attack by antigenic variation and weak immunogenicity. Cell Host and Microbe 10:147–157.

15. Lannergård J, Kristensen BM, Gustafsson MCU, Persson JJ, Norrby-Teglund A, Stålhammar-Carlemalm M, Lindahl G. 2015. Sequence variability is correlated with weak immunogenicity in Streptococcus pyogenes M protein. Microbiologyopen 4:774–789.

16. Metzgar D, Zampolli A. 2011. The M protein of group A streptococcus is a key virulence factor and a clinically relevant strain identification marker. Virulence 2:402–412.

17. Cunningham MW. 2014. Rheumatic Fever, Autoimmunity and Molecular Mimicry: The Streptococcal Connection. Int Rev Immunol 33:314–329.

18. Good MF, Pandey M, Batzloff MR, Tyrrell GJ. 2015. Strategic development of the conserved region of the M protein and other candidates as vaccines to prevent infection with group A streptococci. Expert Review of Vaccines 14:1459–1470.

19. Ochel A, Rohde M, Chhatwal GS, Talay SR. 2014. The M1 Protein of Streptococcus pyogenes Triggers an Innate Uptake Mechanism into Polarized Human Endothelial Cells. Journal of Innate Immunity 6:585–596.

20. Speziale P, Arciola CR, Pietrocola G. 2019. Fibronectin and Its Role in Human Infective Diseases. Cells https://doi.org/10.3390/cells8121516.

21. Cue D, Lam H, Cleary PP. 2001. Genetic dissection of the Streptococcus pyogenes M1 protein: Regions involved in fibronectin binding and intracellular invasion. Microbial Pathogenesis 31:231–242.

22. Henderson B, Nair S, Pallas J, Williams MA. 2011. Fibronectin: A multidomain host adhesin targeted by bacterial fibronectin-binding proteins. FEMS Microbiology Reviews 35:147–200.

23. Hottes AK, Freddolino PL, Khare A, Donnell ZN, Liu JC, Tavazoie S. 2013. Bacterial Adaptation through Loss of Function 9.

24. Price MN, Wetmore KM, Deutschbauer AM, Arkin AP. 2016. A comparison of the costs and benefits of bacterial gene expression. PLoS ONE 11:1–22.

25. Brown EL, Kim JH, Reisenbichler ES, Höök M. 2005. Multicomponent Lyme vaccine: Three is not a crowd. Vaccine 23:3687–3696.

26. El Ghany MA, Jansen A, Clare S, Hall L, Pickard D, Kingsley RA, Dougan G. 2007. Candidate live, attenuated Salmonella enterica serotype typhimurium vaccines with reduced fecal shedding are immunogenic and effective oral vaccines. Infection and Immunity 75:1835–1842.

27. Bahnan W, Happonen L, Khakzad H, Ahnlide VK, de Neergaard T, Wrighton S, Bratanis E, Tang D, Hellmark T, Björck L, Shannon O, Malmström L, Malmström J, Nordenfelt P. 2021. Protection induced by a human monoclonal antibody recognizing two different epitopes in a conserved region of streptococcal M proteins. bioRxiv.

28. Kumra Ahnlide V, Kumra Ahnlide J, Wrighton S, Beech JP, Nordenfelt P. 2022. Nanoscale binding site localization by molecular distance estimation on native cell surfaces using topological image averaging. Elife 11.

29. Macheboeuf P, Buffalo C, Fu C-Y, Zinkernagel AS, Cole JN, Johnson JE, Nizet V, Ghosh P. 2011. Streptococcal M1 protein constructs a pathological host fibrinogen network Atomic coordinates and structure factors for M1 BC1-FgD (2XNX) and M1 A-FgD (2XNY) have been deposited with the Protein Data Bank. HHS Public Access. Nature 472:64–68.

30. Nordenfelt P, Waldemarson S, Linder A, Mörgelin M, Karlsson C, Malmström J, Björck L. 2012. Antibody orientation at bacterial surfaces is related to invasive infection. Journal of Experimental Medicine 209:2367–2381.

31. Frick IM, Wikstrom M, Forsen S, Drakenberg T, Gomi H, Sjobring U, Bjorck L. 1992. Convergent evolution among immunoglobulin G-binding bacterial proteins. Proc Natl Acad Sci U S A 89:8532–8536.

32. Cue D, Dombek PE, Lam H, Cleary PP. 1998. Streptococcus pyogenes serotype M1 encodes multiple pathways for entry into human epithelial cells. Infection and Immunity 66:4593–4601.

33. Talay SR, Valentin-Weigand P, Jerlstrom PG, Timmis KN, Chhatwal GS. 1992. Fibronectin-binding protein of streptococcus pyogenes: Sequence of the binding domain involved in adherence of streptococci to epithelial cells. Infection and Immunity 60:3837–3844.

34. Ferretti JJ, McShan WM, Ajdic D, Savic DJ, Savic G, Lyon K, Primeaux C, Sezate S, Suvorov AN, Kenton S, Lai HS, Lin SP, Qian Y, Jia HG, Najar FZ, Ren Q, Zhu H, Song L, White J, Yuan X, Clifton SW, Roe BA, McLaughlin R. 2001. Complete genome sequence of an M1 strain of streptococcus pyogenes. Proc Natl Acad Sci U S A 98:4658–4663.

35. Nyberg P, Sakai T, Cho KH, Caparon MG, Fässler R, Björck L. 2004. Interactions with fibronectin attenuate the virulence of Streptococcus pyogenes. EMBO Journal 23:2166–2174.

36. Singh K v., Rosa SL la, Somarajan SR, Roh JH, Murray BE. 2015. The fibronectin-binding protein EfbA contributes to pathogenesis and protects against infective Endocarditis caused by Enterococcus faecalis. Infection and Immunity 83:4487–4494.

37. Akerstrom B, Lindahl G, Bjorck L, Lindqvist A. 1992. Protein Arp and protein H from group A streptococci: Ig binding and dimerization are regulated by temperature. J Immunol 148:3238–32343.

38. Nilson BHK, Frick IM, Åkesson P, Forsén S, Bjorck L, Åkerströnv B, Wikstrom M. 1995. Structure and Stability of Protein H and the Ml Protein from Streptococcus pyogenes. Implications for Other Surface Proteins of Gram-Positive Bacteria. Biochemistry 34:13688–13698.

39. Brennan C, Christianson K, Surowy T, Mandecki W. 1994. Modulation of enzyme activity by antibody binding to an alkaline phosphatase-epitope hybrid protein. Protein Engineering, Design and Selection 7:509–514.

40. Accolla RS, Cina R, Montesoro E, Celada F. 1981. Antibody-mediated activation of genetically defective Escherichia coli β-galactosidases by monoclonal antibodies produced by somatic cell hybrids. Proc Natl Acad Sci U S A 78:2478–2482.

41. Klonisch T, Delves PJ, Berger P, Panayotou G, Lapthorn AJ, Isaacs NW, Wick G, Lund T, Roitt IM. 1996. Relative location of epitopes involved in synergistic antibody binding using human chorionic gonadotropin as a model. European Journal of Immunology 26:1897–1905.

42. Massart S, Maiter D, Portetelle D, Adam E, Renaville R, Ketelslegers JM. 1993. Monoclonal antibodies to bovine growth hormone potentiate hormonal activity in vivo by enhancing growth hormone binding to hepatic somatogenic receptors. Journal of Endocrinology 139:383–393.

43. Cunningham MW. 2016. Post-Streptococcal Autoimmune Sequelae: Rheumatic Fever and Beyond. Streptococcus pyogenes: Basic Biology to Clinical Manifestations 1–37.

44. McNitt DH, Choi SJ, Allen JL, Hames RA, Weed SA, van de Water L, Berisio R, Lukomski S. 2019. Adaptation of the group A Streptococcus adhesin Scl1 to bind fibronectin type III repeats within wound-associated extracellular matrix: implications for cancer therapy. Molecular Microbiology 112:800–819.

45. Holden MTG, Scott A, Cherevach I, Chillingworth T, Churcher C, Cronin A, Dowd L, Feltwell T, Hamlin N, Holroyd S, Jagels K, Moule S, Mungall K, Quail MA, Price C, Rabbinowitsch E, Sharp S, Skelton J, Whitehead S, Barrell BG, Kehoe M, Parkhill J. 2007. Complete genome of acute rheumatic fever-associated serotype M5 Streptococcus pyogenes strain manfredo. J Bacteriol 189:1473–1477.

46. Efron B. 1979. Bootstrap Methods: Another Look at the Jackknife. Statistics (Ber) 7:1–26.

