## Supplementary figures for "Group A streptococci induce high-affinity M protein-fibronectin interaction when specific human antibodies are bound"

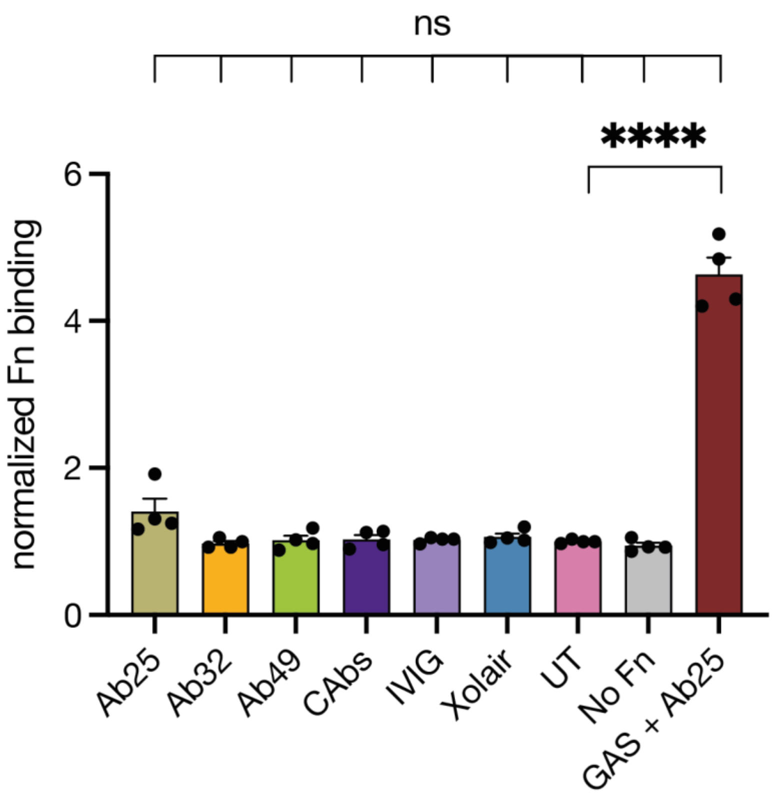


**Supplementary Figure 1** Wells were coated with various antibodies (10µg/ml) not lead to significant binding of Fn (10µg/ml) compared to an untreated control (UT) where wells were not coated with antibodies. Wells coated with GAS and then treated with the same concentration of Ab (GAS + Ab25) did. Error bars represent the SEM. Statistical significance was assessed using one-way ANOVA combined with Dunnett's multiple comparisons test and **** denotes $p < 0.0001$.

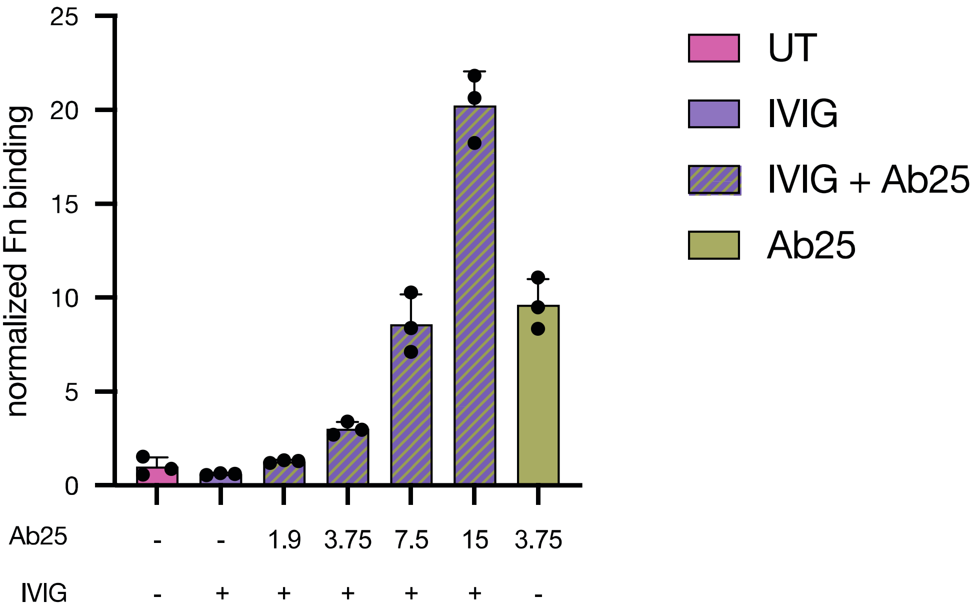


**Supplementary Figure 2** Highly concentrated IVIG (100µg/ml) can be outcompeted by Ab25 (varying concentrations, see graph). All data for this figure was acquired by flow cytometry. Normalized Fn binding signifies the foldchange of median fluorescence intensity compared to GAS only treated with soluble Fn (UT).

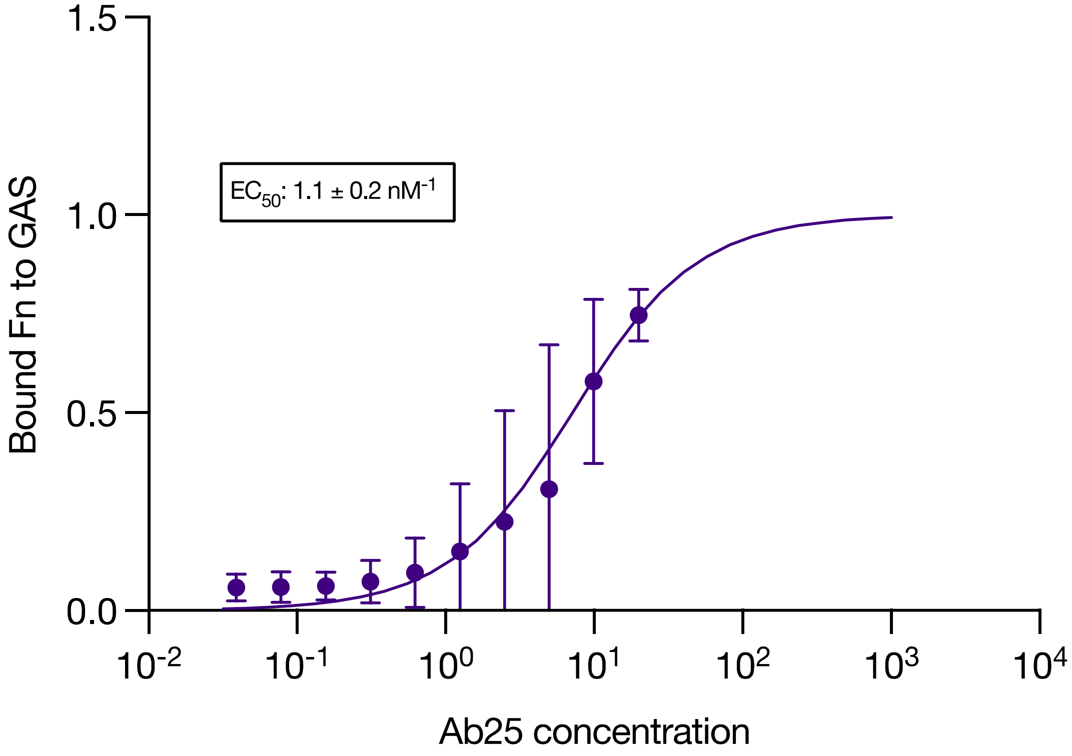


**Supplementary Figure 3** Titration of Ab25 shows positive modulation of fibronectin binding. Fibronectin binding was measured using flow cytometry. The figure shows the measured binding of 1 ug/ml of fibronectin with a fitted ideal binding curve as a function of Ab25 concentration. N=3 for all concentration points. The half maximal effective concentration (EC50) of Ab25 on fibronectin binding is given in the plot together with a confidence interval calculated using the Bootstrap method.

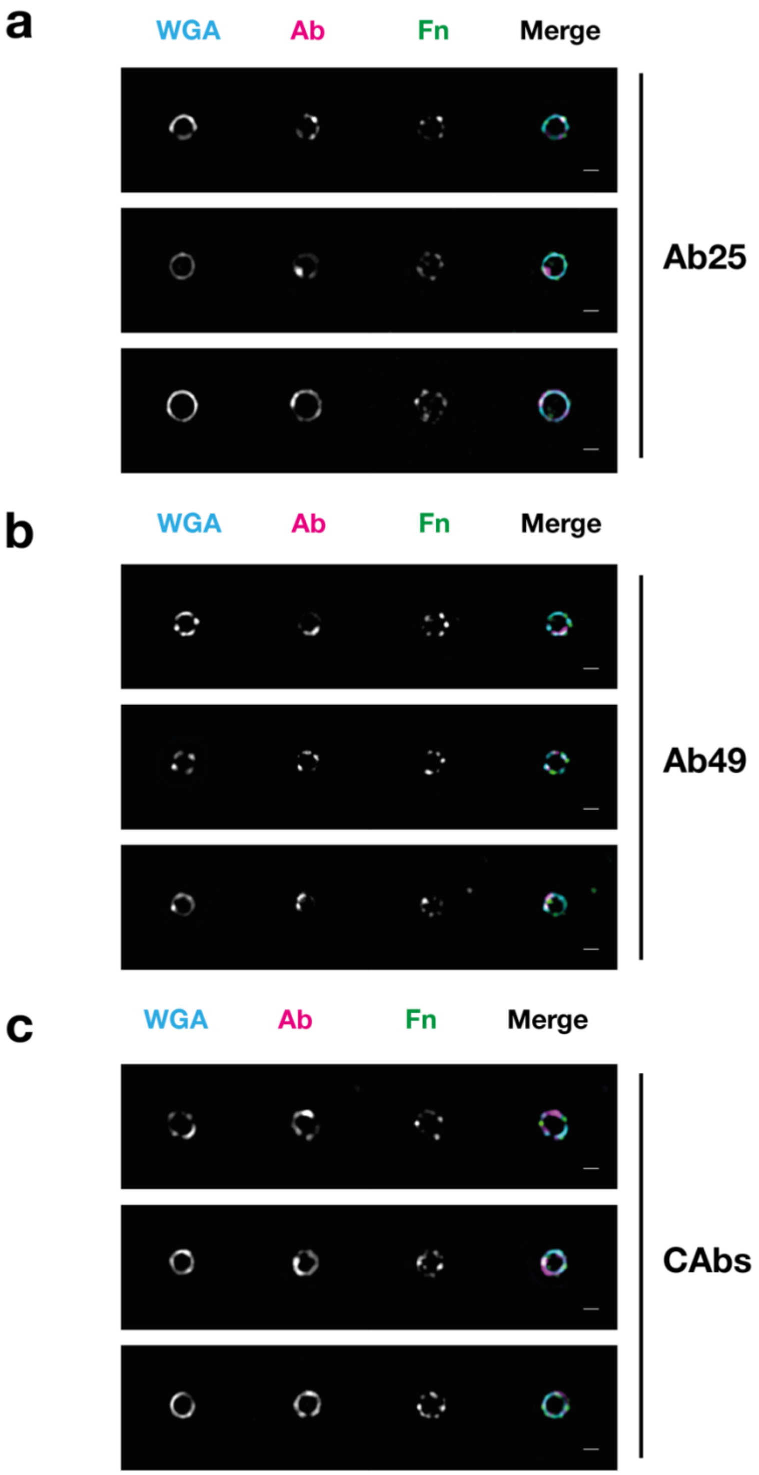


**Supplementary Figure 4** Representative fluorescence colocalization images of SF370 imaged with a N-SIM microscope. The cell wall was stained wheat germ agglutinin (WGA). The bacteria were then treated with Fn and one of 3 antibody treatments. These were **a,** Ab25, **b,** Ab49, and **c,** antibodies derived from a GAS infection convalescent donor (CAbs, bottom). On the right side, merged images of the WGA-stained cell wall, Abs, and Fn are represented by cyan, magenta, and green respectively. The scale bar represents 1µm.
